# A data-driven approach to optimising the encoding for multi-shell diffusion MRI with application to neonatal imaging

**DOI:** 10.1101/661348

**Authors:** J-Donald Tournier, Daan Christiaens, Jana Hutter, Anthony N. Price, Lucilio Cordero-Grande, Emer Hughes, Matteo Bastiani, Stamatios N. Sotiropoulos, Stephen M. Smith, Daniel Rueckert, Serena J. Counsell, A. David Edwards, Joseph V. Hajnal

## Abstract

Diffusion MRI has the potential to provide important information about the connectivity and microstructure of the human brain during normal and abnormal development, non-invasively and in vivo. Recent developments in MRI hardware and reconstruction methods now permit the acquisition of large amounts of data within relatively short scan times. This makes it possible to acquire more informative multi-shell data, with diffusion-sensitisation applied along many directions over multiple *b*-value shells. Such schemes are characterised by the number of shells acquired, and the specific *b*-value and number of directions sampled for each shell. However, there is currently no clear consensus as to how to optimise these parameters. In this work, we propose a means of optimising multi-shell acquisition schemes by estimating the information content of the diffusion MRI signal, and optimising the acquisition parameters for sensitivity to the observed effects, in a manner agnostic to any particular diffusion analysis method that might subsequently be applied to the data. This method was used to design the acquisition scheme for the neonatal diffusion MRI sequence used in the developing Human Connectome Project, which aims to acquire high quality data and make it freely available to the research community. The final protocol selected by the algorithm, and currently in use within the dHCP, consists of *b =* 0, 400, 1000, 2600 s/mm^2^ with 20, 64, 88 & 128 DW directions per shell respectively.

**Highlights:**

- A data driven method is presented to design multi-shell diffusion MRI acquisition schemes (*b*-values and no. directions).
- This method optimises the multi-shell scheme for maximum sensitivity to the information content in the signal.
- When applied in neonates, the data suggest that a *b*=0 + 3 shell strategy is appropriate

## Introduction

Diffusion MRI (dMRI) has been the focus of intense research over the last 20 years, holding great promise for investigation of tissue microstructure due to the technique’s unique sensitivity to the micron-scale diffusion of water. In recent years, there has been increasing interest in the use of so-called multi-shell dMRI, driven to a large extent by improvements in image acquisition and reconstruction methods, which can now provide the amount of data required within clinically feasible scan times. A number of promising methods have been proposed to model data of this nature (e.g. Jensen et al., 2005; Jeurissen et al., 2014; Kaden et al., 2015; Sotiropoulos et al., 2013; White et al., 2013; Zhang et al., 2012).

A common requirement for researchers when setting up a multi-shell dMRI protocol is the determination of optimal imaging parameters, particularly the number of *b*-value shells and the number of diffusion-weighting directions and *b*-value per shell. However, there are many different reconstruction algorithms available, and no clear consensus as to which is best. This makes it very difficult to identify optimal imaging parameters, since any optimisation will typically aim to reduce errors in the estimated parameters of a given model; this obviously does not provide any guarantees that the data would be suitable for a different (possibly yet to be devised) reconstruction algorithm. To illustrate the difficulties with such an approach, Figure 1 shows the median absolute deviation in DTI metrics computed using different subsets of an extended data acquisition to seek protocols that produce low errors. The approach is to calculate mean diffusivity (MD) and fractional anisotropy (FA) using a target analysis method of choice for all the available data (6 shells, 50 directions on each, 5 subjects) to determine a “reference value” and then to do the same using only 3 of the shells to find feasible duration protocols that are reliable in the sense of producing the lowest median error (see Appendix A for full details). This was performed using three different commonly-used tensor estimation strategies: ordinary least-squares (OLS), weighted least-squares (WLS), and iteratively reweighted least-squares (IWLS) (Basser, 1995; Koay et al., 2006; Salvador et al., 2005; Veraart et al., 2013), all implemented within *MRtrix3*. Depending on the analysis method used, different “optimal” protocols are identified – see figure caption for details. It is striking that this is the case even though the same model of diffusion is assumed (the diffusion tensor) for all cases.

**Figure 1:**
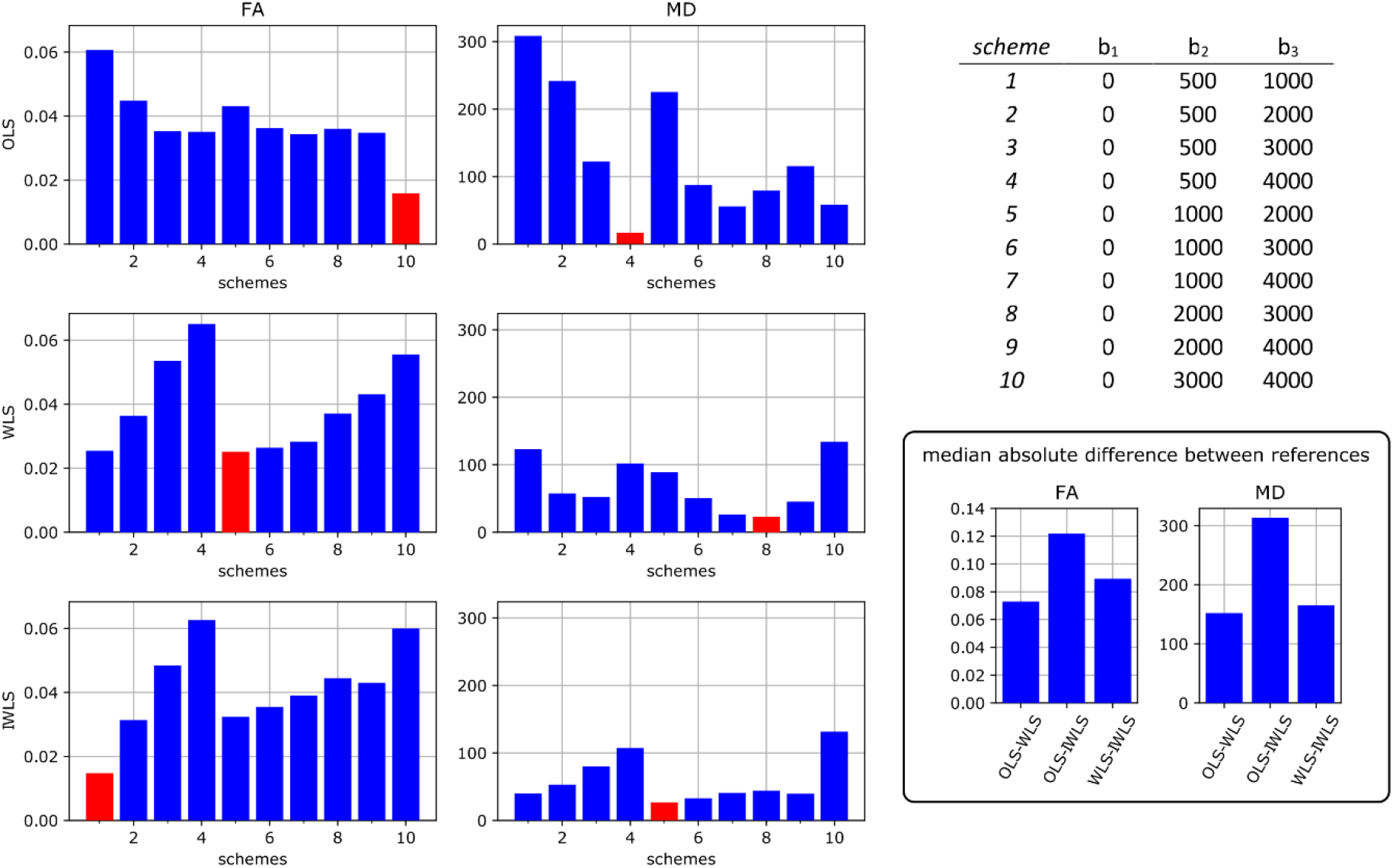
Optimising sampling schemes for measuring FA and MD using minimal discrepancy relative to a superset of all available data for 3 different DTI analysis methods (ordinary least squares (OLS), weighted least squares (WLS) and iteratively reweighted least squares (IWLS). The optimal protocols, which are different in each case, are highlighted in red. The bar graphs show the median absolute difference in DTI metrics obtained using all the data compared to those obtained using different subsets of 3 of the available b-values. The results for MD are shown in units of µm^2^/s (FA is dimensionless). Differences were computed voxel-wise, and the median absolute difference was calculated over the whole brain for each of the 5 subjects included in the study, then averaged across subjects. Results are shown for fractional anisotropy (FA, left) and mean diffusivity (MD, centre), computed using three commonly used estimators: ordinary least-squares (OLS, top row); weighed least-squares (WLS, middle row); and iteratively re-weighted least-squares (IWLS, bottom row). The x-axis corresponds to the numbered schemes shown in the table on the top right, which specify the included shells. The schemes with the lowest discrepancy are highlighted in red. The bottom right panel shows the equivalent median absolute difference between the metrics obtained with the different fitting approaches using all the available data (i.e. the difference between the references used for the other plots).

There is therefore a need for a means to determine acquisition parameters that provide the most eloquent, information-rich data in a manner agnostic to any particular reconstruction algorithm. This is particularly important for large-scale data collection projects that will provide a shared resource to be used by diverse academic groups for purposes that the users rather than the data collectors define.

This study was motivated by the need to optimise the dMRI acquisition scheme for the developing Human Connectome Project (dHCP), which aims to acquire structural, functional and diffusion MRI data from over 1,000 neonates and fetuses, amongst other clinical, behavioural and genetic measures (http://www.developingconnectome.org/). Since these data are to be made freely available to the wider neuroimaging community, it was important to acquire data suitable for the widest possible range of analysis methods. Moreover, the diffusion-weighted (DW) signal in the perinatal age range differs markedly from that in adults, in that it exhibits much higher apparent diffusion (suggesting increased extracellular and reduced intracellular content), and much lower anisotropy. Furthermore, these characteristics vary strongly as a function of both location and age, as the various structures in the brain mature at different rates. Much analysis work to date has used algorithms designed for adult studies, but there is clearly scope for optimising all aspects of the analysis/modelling pipeline and this could result in methods that are different from existing approaches.

Taken together, these factors provide strong motivation to develop a data driven strategy that can be used to design data acquisitions for a desired target subject group, optimised for information content rather than to support a predefined processing method. To address this challenge, we propose to use an information-theoretic approach, similar in spirit to our previous work on optimising single-shell dMRI parameters (Tournier et al., 2013).

## Methods

In this paper, we propose a framework to determine the optimal number of shells, along with the corresponding *b*-value and number of directions sampled. To make this problem tractable, we separate the *b*-value dependence from the orientation dependence and treat each problem separately. For the *b*-value dependence, the central concept is then to identify a linear basis to represent the *b*-value dependence of the data (voxel values) as measured empirically, and determine the number of coefficients of that basis that can realistically be measured in practice. Given this number, the task is then to identify a set of parameters (i.e. number of shells, *b*-value per shell and number of DW volumes per shell) that provide optimal sensitivity to these coefficients.

The angular dependence was itself the focus of previous work, applied to the adult case (Tournier et al., 2013); here, we simply deploy the same methodology to determine the per-shell sampling density required to capture the detectable number of spherical harmonics. Briefly, the approach expresses the dMRI signal for a single *b*-value shell using the spherical harmonics basis, and identifies the highest angular frequency term that can realistically be detected in the data. This analysis is performed only in voxels deemed to contain a single-fibre population, since these contain the highest angular frequency content, and also because this allows averaging across voxels (after realignment of the fibre direction to a common axis). For further details, the reader is referred to (Tournier et al., 2013). This analysis provides an estimate of the angular frequency content at each *b*-value, which translates directly into the corresponding minimum number of directions required for non-lossy sampling (i.e. without aliasing in the angular domain). Once the optimal *b*-values have been identified, this analysis can be used to ensure that the sampling density within each shell is adequate.

To determine the sampling requirements across *b*-values, we focus on the *b*-value dependence of the orientationally-averaged raw DW signal as a function of *b*-value. Assuming sufficiently dense and uniform sampling, the mean DW signal over a single *b*-value shell is rotationally invariant, and independent of the fibre arrangements (Kaden et al., 2015; Raffelt et al., 2012; Reisert et al., 2017). Near-uniform distribution of gradient directions can be obtained using electrostatic repulsion approaches (Jones et al., 1999; Papadakis et al., 2000), as used in this study, and sufficient sampling density can be verified using the approach described above (Tournier et al., 2013).

As an overview, this study consists of the following steps:

- Acquire data sufficient densely sampled in the *b*-value and angular domains to capture all the expected features of the DW signal;
- Identify a suitable data-driven linear basis to represent the signal;
- Estimate the effect size observed in the signal for each coefficient of that basis;
- Identify the set of acquisition parameters that provide optimal sensitivity to these coefficients.

Each of these steps is described in more detail in the following sections.

### Data acquisition and pre-processing

Data were acquired from 5 neonates scanned at term-equivalent age (see Table 1 for details) on a Philips Achieva 3T magnet at the Evelina Newborn Imaging Center at St. Thomas’ Hospital (London, UK). The study was approved by the National Research Ethics Committee and written informed parental consent was obtained prior to scanning. The system was equipped with a standard 32-channel head coil, using a diffusion-weighted pulsed gradient spin-echo EPI sequence (TE/TR = 70/2260 ms, 2×2mm voxel size, 11 slices with 2mm thickness with a 1mm slice gap, 112×112 matrix, SENSE factor 2, single anterior-posterior phase-encode direction). Data were collected along 50 non-collinear directions optimised using electrostatic repulsion (Jones et al., 1999), at *b*-values = 500, 1000, 2000, 3000, 4000 s/mm^2^, along with 5 *b* = 0 s/mm^2^ volumes. These directions were optimised using the *MRtrix3* ‘dirgen’ command, and the same set of 50 directions was used for each shell. We note that in adults, a spherical harmonic order of 8 (45 parameters) was shown to be sufficient to capture all realistically measurable components of the signal, even up to *b* = 5000 s/mm^2^ (Tournier et al., 2013); the 50 directions used in the acquisition here are therefore sufficient to satisfy this criterion, especially considering that neonatal data will exhibit lower anisotropy than observed in adults (Dubois et al., 2008; Hüppi and Dubois, 2006; Yoshida et al., 2013).

**Table 1:** details of the subjects included in this study.

| GA<br>(weeks) | PMA<br>(weeks) | Clinical features | Imaging findings |
| --- | --- | --- | --- |
| 39+2 | 45+4 | Smith-Lemli-Opitz syndrome | Cerebellar hypoplasia, small hypothalamic hamartoma |
| 38+0 | 40+1 | Fetal alcohol syndrome | Small but otherwise normal imaging appearance |
| 40+5 | 42+5 | Seizures, antenatal ventriculomegaly | Perisylvian polymicrogyria |
| 33+3 | 36+1 | Preterm | small germinal matrix haemorrhage |
| 39+6 | 40+5 | Meconium aspiration, mild hypoxia | Normal imaging appearance |

For each subject, the data were first corrected for motion and eddy-current induced distortions using the FSL EDDY (version 5.0.8) (Andersson and Sotiropoulos, 2016, 2015). The mean DW signal for each shell was then computed within a brain mask. Typical images of the resulting mean signal are shown in Figure 2. Analyses were performed both for each subject individually, and for all their data combined (by concatenation along the slice axis). Unless otherwise stated, results shown correspond to the combined data.

**Figure 2:**
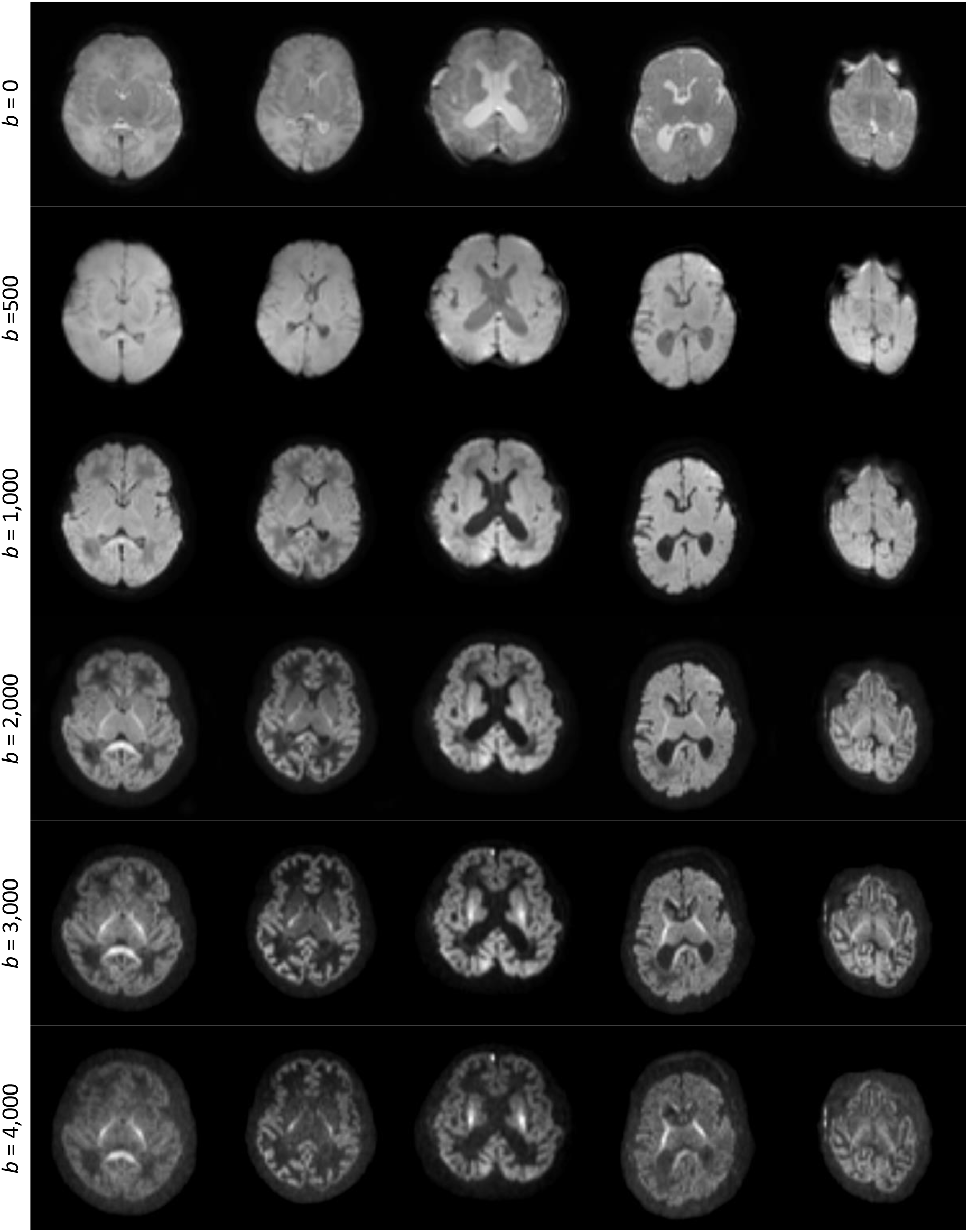
images of the mean raw dMRI signal for all 5 subjects (columns), for each *b*-value (rows). These constitute the input for the optimisation algorithm. Note that the images are windowed independently for each *b*-value to allow visualisation of the contrast in the high *b*-value images.

### Identifying a suitable linear basis

In contrast to the angular domain where spherical harmonics form a natural basis (Tournier et al., 2013), there is no such natural basis to represent the *b*-value dependence of the signal. While some bases have been proposed (Assemlal et al., 2009; Caruyer and Deriche, 2012; Descoteaux et al., 2011; Fick et al., 2016; Hosseinbor et al., 2013; Merlet and Deriche, 2013; Özarslan et al., 2013; Rathi et al., 2014), there is an inherent scale dependence that makes the analysis dependent on user-defined parameters (Christiaens et al., 2019). For this reason, we adopted a fully-data driven, model-free basis, derived using matrix decomposition approaches.

We use the compact SVD decomposition, which decomposes a general *m* × *n* matrix *A* (with *m* ≤ *n*) as *USV*^*T*^, where *U* is a *m* × *m* orthonormal matrix, *S* is a *m* × *m* diagonal matrix of singular values, and *V*^*T*^ is a *m* × *n* orthonormal matrix. To make use of this decomposition, we form the compact SVD of the transpose of *D*:

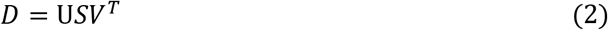

More specifically, we consider a multi-shell protocol consisting of *N*_*s*_ *b*-values, with *n*_*s*_ uniformly distributed DW directions for shell index *s*. We form the *N*_*s*_ × *N*_*v*_ matrix *D* consisting of the mean DW signal for each brain voxel at each *b*-value, where *N*_*v*_ is the number of voxels in the subject-specific brain mask. We wish to decompose *D* into a *N*_*s*_ × *N*_*v*_ matrix *W* of per-voxel weights, and a *N*_*s*_ × *N*_*s*_ orthonormal basis matrix *H*, such that:

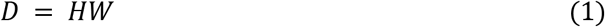

The weight and basis matrices can then be obtained as:

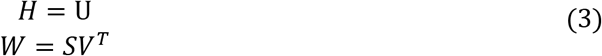

Furthermore, thanks to the properties of the SVD, the singular values (i.e. the diagonal of *S*) can be used to obtain an estimate of the typical component-wise effect sizes *ϵ* (this evaluates to the root-mean-square value of the weights across the rows of *W*):

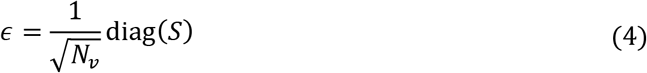

### Sensitivity analysis

The decomposition described above provides a linear breakdown of the effects observed in the data, and of their relative contributions. This can be used to predict the sensitivity to these various effects for a candidate acquisition with the same *b*-value shells, but different numbers of volumes per shell. Assuming independent and identically distributed (i.i.d.) Gaussian noise, the expected variance in the measured mean DW signal for each shell *s* is simply 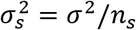, with *n*_*s*_ the number of volumes in shell *s*, and *σ*^2^ the variance per measurement in the data. This rests on the observation that provided the sampling density is uniform and sufficiently dense (as per (Tournier et al., 2013)), the variance in the mean DW signal for a given shell is inversely proportional to the number of DW volumes acquired at that *b*-value; the mean DW signal is orientationally invariant, and each volume contributes equally to its estimation.

The predicted variance in the mean per-shell signal can be used to predict the variance in the estimated basis coefficients using the law of propagation of errors (Arras, 1998). Based on equation (1), if *d* = *Hw* is the vector of mean per-shell signals for a single voxel (where *w* is the corresponding vector of basis coefficients), the variance-covariance matrix Σ_*w*_ for the coefficients *w* is related to the corresponding variance-covariance matrix Σ_*d*_ for the mean per-shell signals *d* according to:

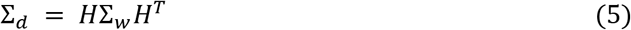

Since *H* is orthogonal by construction, *H*^−1^ = *H*^*T*^, yielding:

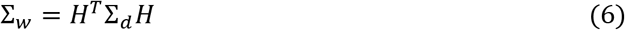

With repeated measurements per shell, 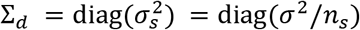 (see above). Hence:

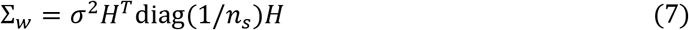

Finally, we compute the coefficient of variation (CV) for coefficient *c* as the ratio of its standard deviation (given by the square-root of its variance Σ_*w*_(*c*, *c*)) to its effect size *ϵ*_*c*_:

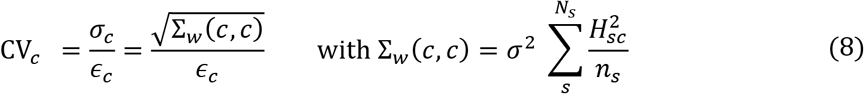

Note that CV_*c*_ is equivalent to the inverse of the contrast to noise ratio (CNR) of the coefficient of interest.

### Optimising the number of directions per shell

The optimal acquisition should strive to minimise the combined coefficient of variation of the *N*_*c*_ estimated coefficients. We therefore use the *sum of squared coefficients of variation* (SSCV) as a measure of optimality:

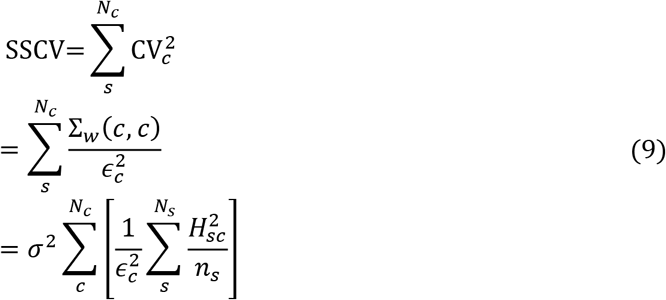

Note that if the effect size were the same for all coefficients, this is proportional to the sum of the variances, which is appropriate for independent measurements. However, when effect sizes differ, this measure is most strongly influenced by the coefficients with the highest coefficient of variation, pushing the optimisation towards parameters that maximise the CNR of the noisiest components, without overly penalising high CNR components.

We now need to derive the number of directions per shell ***n*** = {*n*_*s*_} that maximise the sensitivity, and hence minimise the SSCV, within the constraint that the number of DW volumes, *N*_total_, is fixed by scan time limitations:

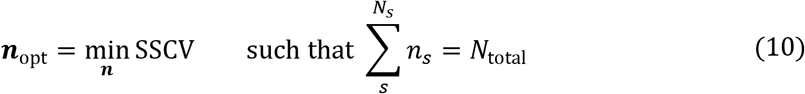

This can be solved using Lagrangian multipliers:

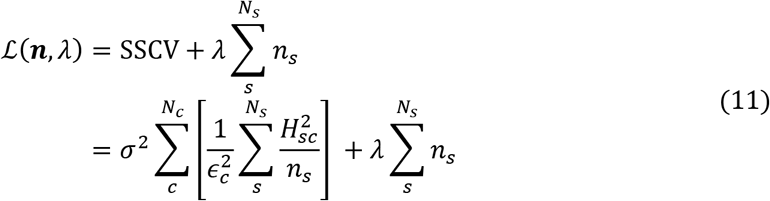

The derivative with respect to the number of directions in shell *s* is:

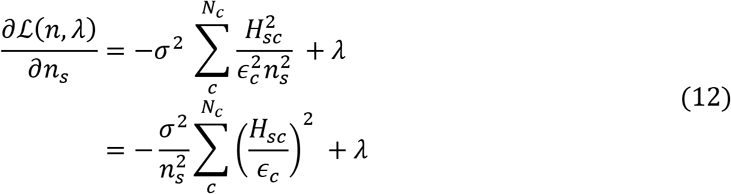

Setting the derivative to zero yields:

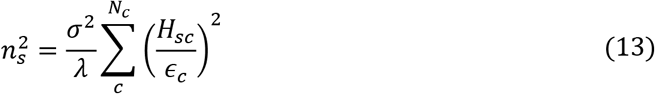

This therefore provides the optimal number of DW volumes for shell *s relative* to the other shells. Since we assume that the total number of volumes is fixed by scan time constraints, the absolute numbers can trivially be obtained by normalising to *N*_total_:

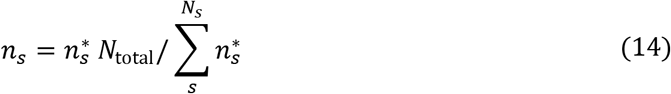

where 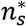 is the value obtained by setting *λ* in equation (14) above to an arbitrary constant. It is interesting to note that with this derivation, the fractions of measurements on each shell is independent of *N*_total_ or *σ*: they depend purely on the intrinsic effects in the DW signal.

### Generalisation to arbitrary *b*-values

So far, the derivation above holds for data acquired over a fixed number of shells with pre-determined *b*-values. If a sufficiently large number of different combinations of shells and *b*-values were available, this could be used to identify the optimal set of parameters, by searching for the subset of *b*-values with minimum SSCV (equation (9)). It is however impractical to acquire data over a sufficient number of *b*-values to ensure the search is exhaustive, particularly in a neonatal cohort where scan times must be kept short.

We therefore used interpolation methods to resample the acquired data, allowing the generation of *predicted* data for any combination of *b*-values; this approximation is justified on the grounds that the diffusion signal varies smoothly as a function of *b*-value, so that interpolation errors are likely to be minimal. For this purpose, we use the *Piecewise cubic Hermite Interpolating Polynomial* (PCHIP) algorithm (Higham, 1992) available in Matlab (The MathWorks Inc, Natick, Massachusetts), which has the desirable property of explicitly preserving monotonic behaviour.

Given a fixed desired number of shells, the optimal *b*-values for each shell are determined using a simple non-linear optimisation approach (the Nelder-Mead simplex algorithm (Nelder and Mead, 1965), as implemented in Matlab’s *fminsearch* routine). For each iteration, the mean DW signals per voxel at the current *b*-values are obtained by PCHIP interpolation from the measured data. The SSCV of the parameters is then optimised with respect to the number of directions per shell as described in the section *Optimising the number of directions per shell* above. Finally, the optimised SSCV value is used as the cost function for the search over *b*-values.

### Accounting for *T*_2_ decay

So far, the derivation has ignored the effect of *T*_2_ decay on the sensitivity of the acquisition. In practice, the shell with the highest *b*-value will dictate the echo time of the entire acquisition (assuming a constant echo time is desired, as is typically the case), and this will have a direct impact on the SNR of the data. To account for these effects, we include a *b*_max_-dependent *T*_2_ decay term, assuming a standard pulsed gradient spin-echo (PGSE) sequence, similar to (Alexander and Barker, 2005), with square diffusion-sensitising gradients and the following parameters:

- a 5ms delay between the 90° excitation pulse and the onset of the first DW gradient pulse;
- a 5ms pulse duration for the 180° refocusing pulse (including slice selection gradients and crushers, if any);
- a 15ms delay between the end of the second DW gradient pulse and echo time;
- 80 mT/m gradient strength.

The attenuation factor is then computed as exp(−*TE*/*T*_2_), and is used to scale the effect sizes in equation (4), inherently penalising parameters sets that include a high *b*-value shell. Results were generated assuming *T*_2_ = 100, 150 & 200 ms, which are typical for neonatal brain (Williams et al., 2005), as well as ignoring *T*_2_ relaxation.

## Results

The angular domain analysis shows that the angular frequency content is much lower in neonates (Figure 3) than has been shown previously in adults (Tournier et al., 2013), as expected. In our data, spherical harmonic terms of order 8 and above were not detectable. Order 6 terms were best observed with *b* ≥ 2000 s/mm^2^, and imply that a minimum of 28 directions are required at these *b*-values. At lower *b*-values, only SH terms of order 2 or 4 could be detected, implying a minimum of 6 and 15 directions respectively.

**Figure 3:**
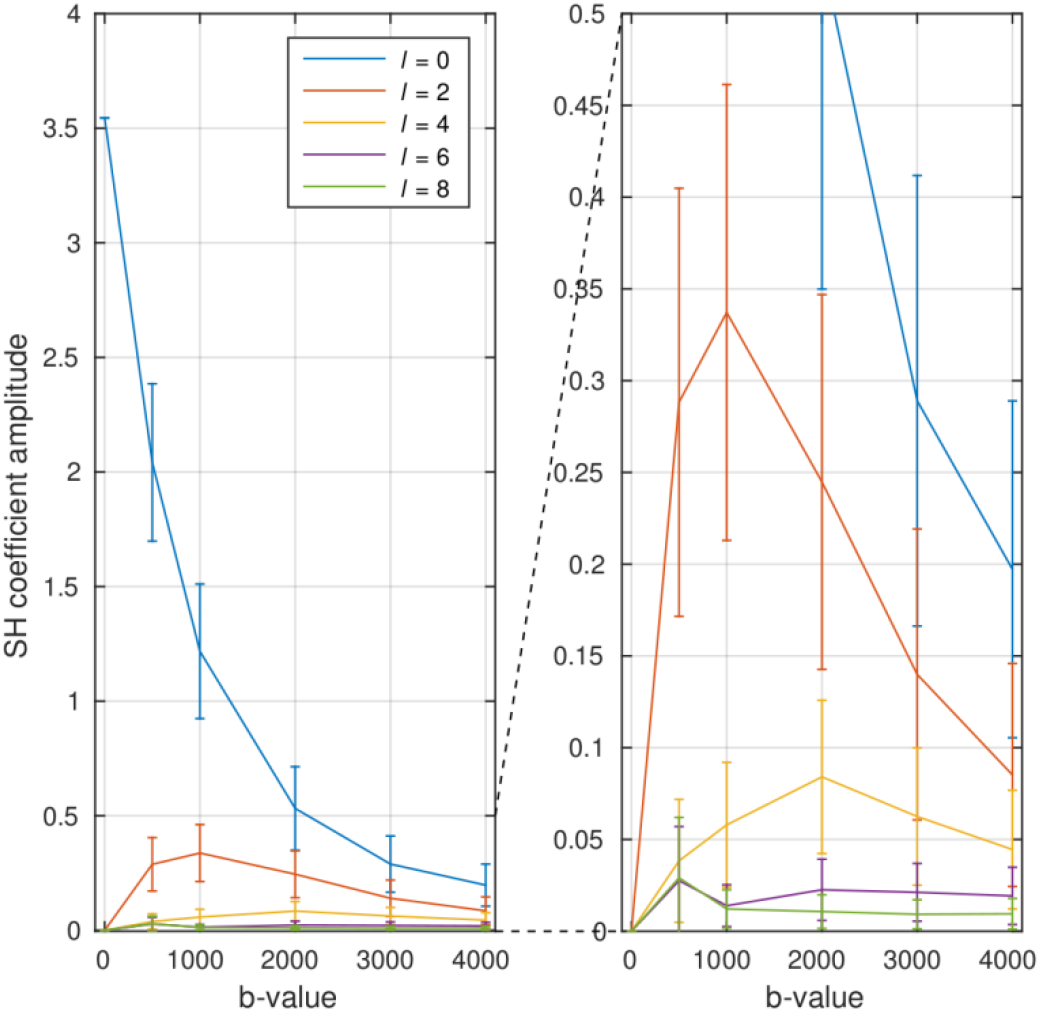
angular frequency content of dMRI signal in neonates, as a function of *b*-value (cf. (Tournier et al., 2013)). Top: polar plot of the signal (fibre orientation is along vertical axis), showing the rapid decrease in signal with increasing *b*-value, along with the expected increase in angular contrast. Bottom: a plot of the corresponding spherical harmonic coefficient amplitudes. Angular frequencies above *l* = 4 are already within the noise floor, although there is a suggestion that *l* = 6 might be detectible for *b* ≥ 2,000 s/mm^2^. This implies that a minimum of 15 & 28 DW directions are required for *b* < 2000 s/mm^2^ and *b* ≥ 2,000 s/mm^2^ respectively.

The responses derived using SVD are shown in Figure 4, along with their corresponding weights maps in Figure 5, and are remarkably consistent across subjects. The first component corresponds to the mean DW signal over all voxels, with the higher-order components showing increasingly rapid oscillations with b-value. Clear structure can be observed in the corresponding weights maps for at least the first 4 components (although the 4th component may possibly contain some residual misregistration due to e.g. motion or eddy-currents, as well as any actual biology related structure). In our data, the 5th order lacks consistency across subjects, suggesting that the remaining spatial structure may be strongly related to artefacts in the data. The 6^th^ and highest order that could be derived from the test data given the number of shells, lacks both consistency across subjects and anatomical coherence, suggesting that noise and artefacts have become completely dominant. The effect sizes for these components decrease rapidly for the higher-order coefficients, as shown in Table 2.

**Table 2:** effect sizes for each observed component in the decomposition (as shown in Figure 4, across all subjects (arbitrary units).

| component | 1 | 2 | 3 | 4 | 5 | 6 |
| --- | --- | --- | --- | --- | --- | --- |
| effect size | 1.000 | 0.134 | 0.038 | 0.018 | 0.009 | 0.004 |

**Figure 4:**
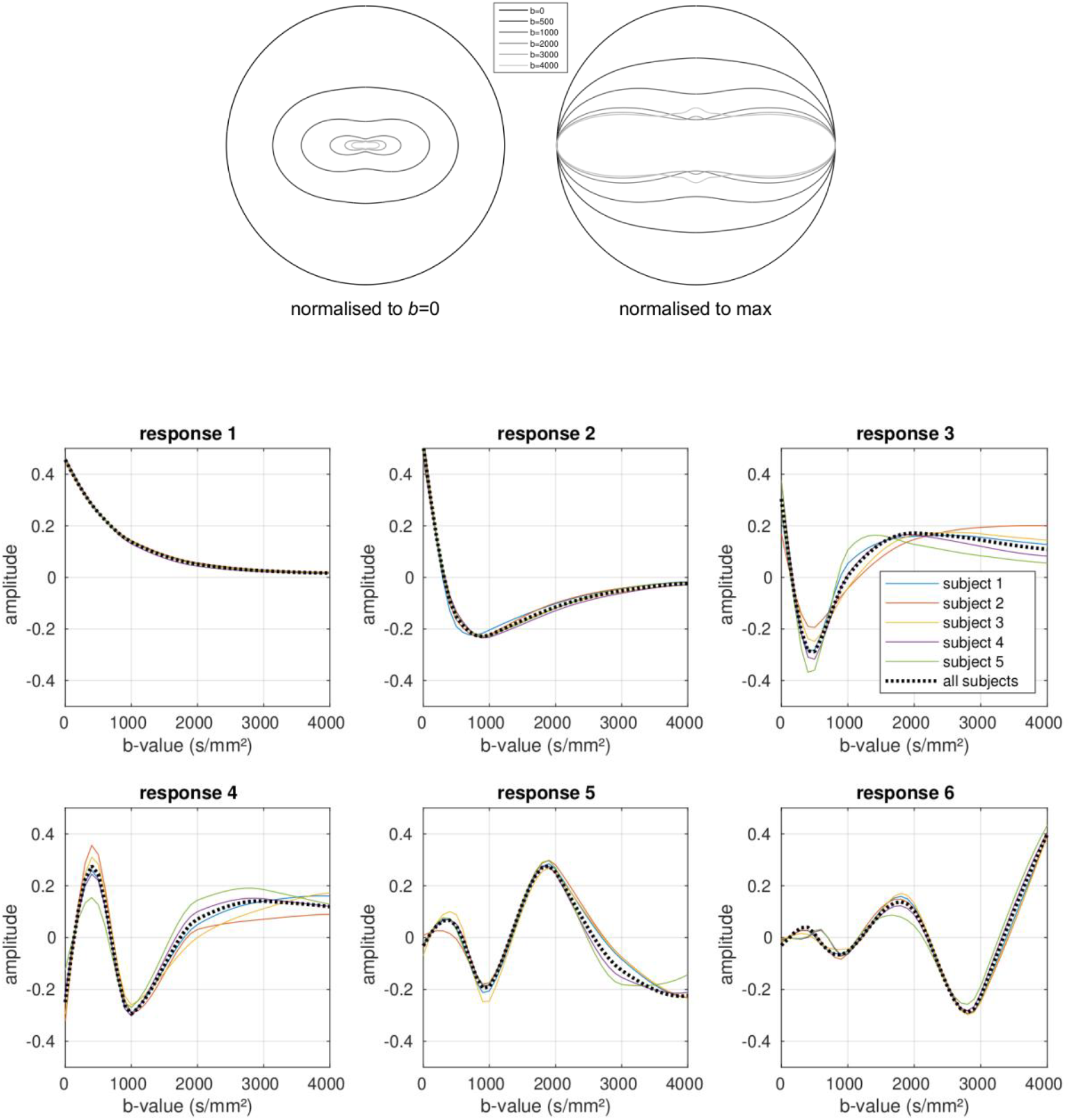
the responses observed in the data, for all 5 subjects included in the study. The responses estimated from each subject independently are shown by the solid coloured lines. The dotted black line shows the responses estimated from the combined data across all subjects. See Table 2 for the corresponding effect sizes.

**Figure 5:**
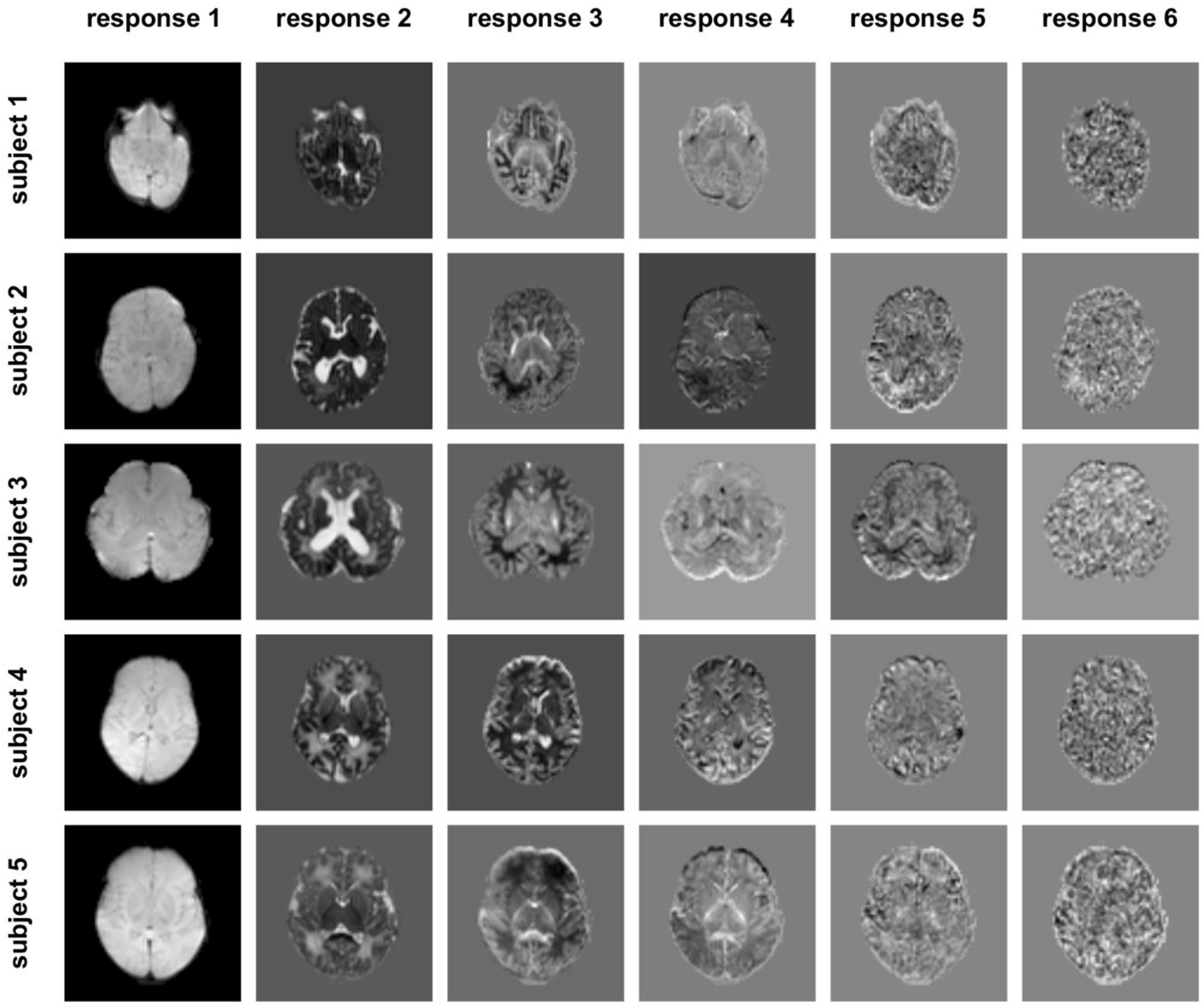
weights maps for each component in Figure 2, for all 5 subjects included in the study.

The optimal *b*-values selected by the algorithm are shown in Figure 6 for the case of 2, 3 & 4 shells (not including the *b*=0 shell), assuming a *T*_2_ value of 150 ms in neonatal brain tissue (Williams et al., 2005). Figure 7 shows the effect of different *T*_2_ values for the 3 shell case. In all cases, a low *b*-value in the region of *b* = 400 s/mm^2^ is included, and can be seen to provide good separation between the first component and the other smaller components. In general, the algorithm seems to distribute *b*-values in a way that provides optimal separation between the different components, as desired. When more shells are included, the highest *b*-value selected increases, particularly when *T*_2_ relaxation is ignored. When relaxation effects are included, the maximum *b*-value never exceeds *b*≈2,600 s/mm^2^.

**Figure 6:**
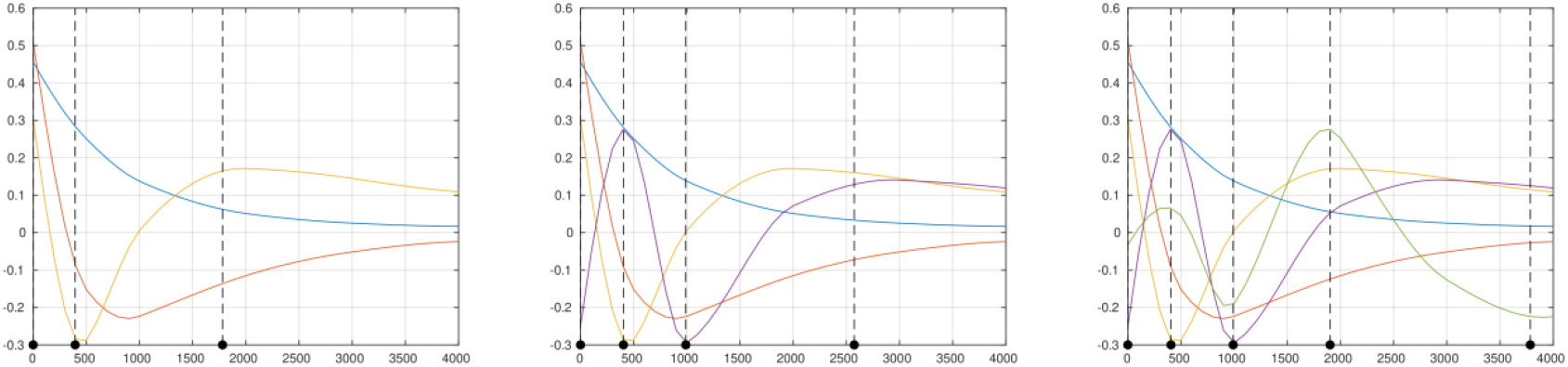
optimal *b*-values (dotted lines) selected by the algorithm, for 2-shell (left), 3-shell (middle) and 4-shell (right) cases, overlaid on the corresponding basis functions. Results obtained assuming a *T*_2_ value of 150 ms.

**Figure 7:**
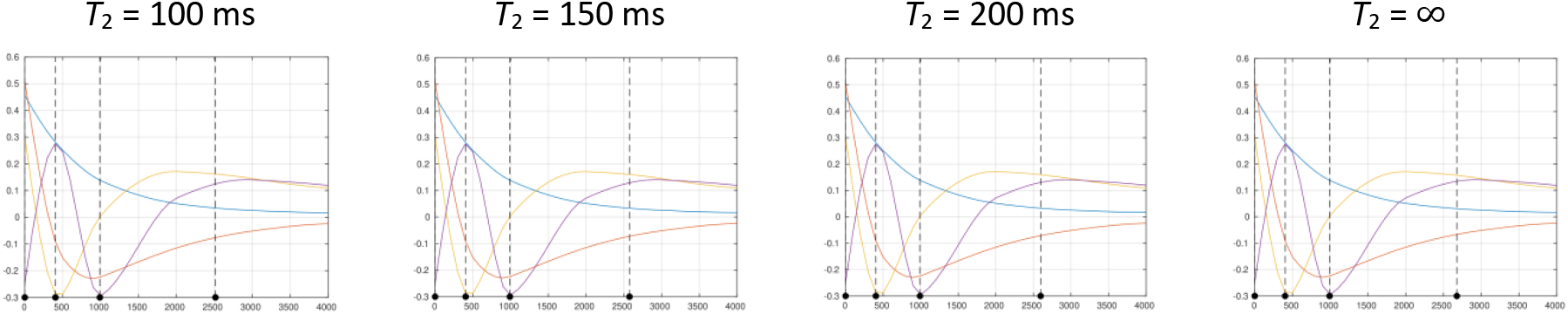
effect of *T*_2_ on optimal *b*-values selected, for the 3-shell case. While *T*_2_ does have an influence, it is relatively minor, with lower *T*_2_ leading primarily to a slightly reduced maximum *b*-value, with little to no impact on the other selected *b*-values.

The *b*-values selected as optimal are stable across subjects, apart from the maximum *b*-value, as shown in Figure 8 for the 3-shell case. The maximum *b*-value varies in this case from *b* ≈ 2,300 to 3,500 s/mm^2^; this is likely related to the signal levelling out as a function of *b*-value in this range, leading to a broad optimum.

**Figure 8:**
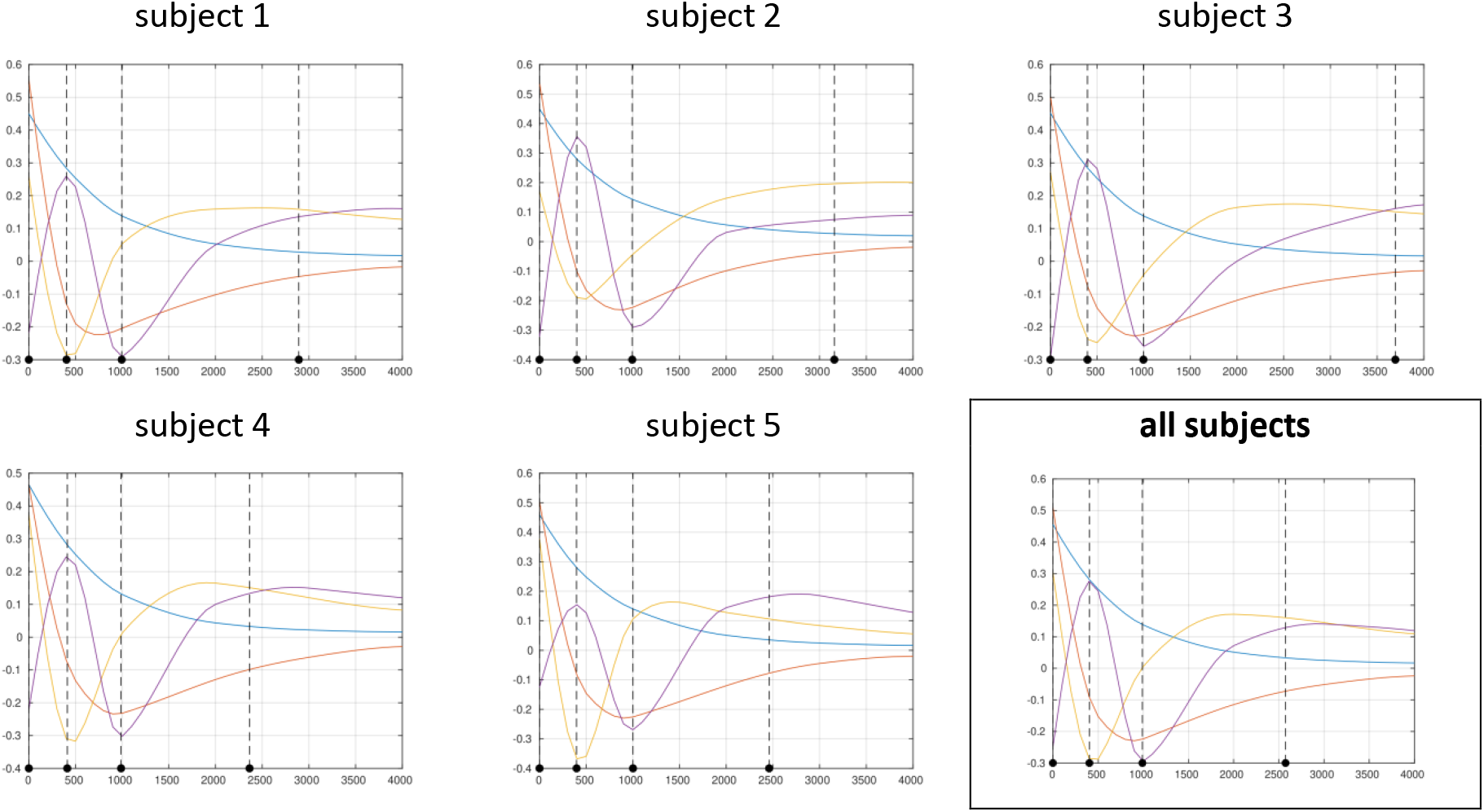
optimal *b*-values selected by the algorithm when applied to each subject individually, for the 3-shell case and assuming *T*_2_ = 150 ms. While there is some variation in the maximum *b*-value, the other *b*-values remain very stable.

Table 3 shows the optimal results for 2, 3 & 4 shells assuming *T*_2_ = 150 ms (suitable for neonates at 3T (Williams et al., 2005)). Along with the optimal *b*-values, the optimal number of directions per shell are also listed as a percentage of the total number of volumes acquired. For reference, the contrast-to-noise ratio (CNR) for the coefficients themselves is shown in Table 4, assuming SNR = 30, and *N*_*total*_ = 100 imaging volumes. As expected, the CNR drops rapidly for the smaller coefficients, and is typically improved by using the smallest number of shells needed to characterise that coefficient (i.e. *b*=0 + 2 shells to characterise the 3rd coefficient). These CNR calculations can be generalised to arbitrary conditions by adjusting them based on the actual number of volumes and SNR_*b*=0_ values:

**Table 3:** optimal *b*-values (in s/mm^2^) for each shell and corresponding number of directions (*N*_*total*_), expressed as a percentage of the total number of volumes acquired. Results are shown for the 2 shell, 3 shell and 4 shell cases, assuming a *T*_2_ value of 150 ms.

|  |  |  |  |  |  |  |
| --- | --- | --- | --- | --- | --- | --- |
| 2 shells | $b$-value | 0 | 395 | 1,781 | | |
| | $N_{total}$ | 12% | 29% | 59% | | |
| 3 shells | $b$-value | 0 | 404 | 990 | 2,574 | |
| | $N_{total}$ | 6% | 19% | 28% | 47% | |
| 4 shells | $b$-value | 0 | 405 | 990 | 1,901 | 3,784 |
| | $N_{total}$ | 3% | 9% | 16% | 29% | 43% |

**Table 4:** contrast to noise ratios (CNR) for each component, as would be estimated using the corresponding parameters in Table 3. These values assume a *T*_2_ value of 150 ms, a total of *N*_*total*_ = 100, and an SNR in the *b*=0 images of 30.

| component | 1 | 2 | 3 | 4 | 5 |
| --- | --- | --- | --- | --- | --- |
| **2 shells** | 152 | 18 | 6.0 |  |  |
| **3 shells** | 125 | 16 | 4.5 | 2.7 |  |
| **4 shells** | 88 | 13 | 4.0 | 2.1 | 1.3 |

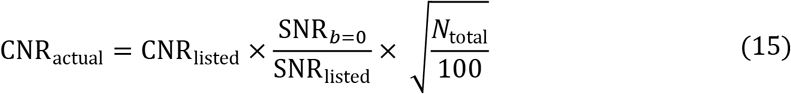

As expected, larger numbers of volumes are needed for the higher *b*-values shells. This is expected given the lower SNR available at these *b*-values, and also helps to ensure that the minimum orientation sampling density requirements (identified earlier using the angular frequency analysis) are satisfied (Tournier et al., 2013).

Based on this optimisation a 3 shell protocol was adopted for the dHCP (Hutter et al., 2017). The acquisition includes a total of 300 samples distributed in the proportions listed in table 3. Figure 9 shows example images and shell average images acquired as part of the dHCP study (PGSE EPI acquisition with multiband factor 4, 64 slices, TE/TR = 90/3800 ms, 1.5×1.5×3mm resolution, 4 phase-encoding directions, SENSE 1.2, Partial Fourier 0.855), after correction for EPI distortion, subject motion and outliers (Andersson and Sotiropoulos, 2016, 2015). While the individual images in the highest shell display the expected low SNR, the shell average reveals highly coherent anatomical structure confirming its information richness, which can be well accessed by virtue of the large number of samples collected. All shells have angular sampling in excess of the requirements determined from the pilot data.

**Figure 9:**
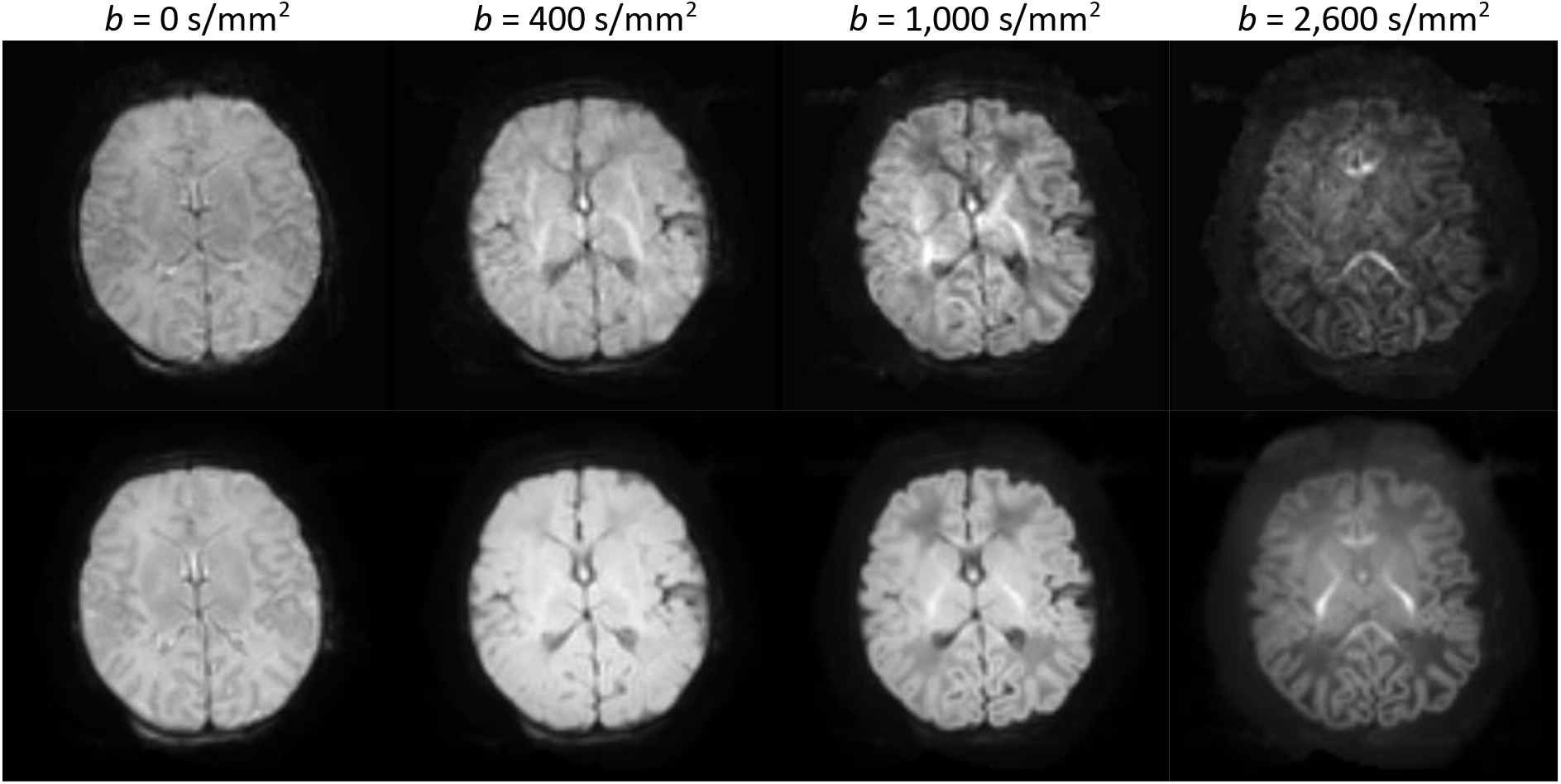
single diffusion-weighted images (top row) and mean diffusion-weighted image (bottom row) for each shell for a typical subject acquired using the final dHCP protocol.

## Discussion

This study provides a data-driven framework to identify the optimal parameters for multi-shell HARDI acquisitions, based on an information-theoretic approach. It is independent of any particular reconstruction/diffusion analysis algorithm, which is a significant advantage given that many of these approaches are under active development, have a number of different tuneable parameters, and are designed to provide different types of information. As illustrated in Appendix A, a protocol optimised for any one of these would therefore not be guaranteed optimal for any of the other methods (or indeed, any other parameters or outputs of the same method). The proposed design method allows protocols to be implemented now that will remain future-proof as novel analysis approaches are developed.

While a number of decompositions could be used for the *b*-value dependence, singular value decomposition is arguably the most appropriate for this study. The SVD provides a unique and complete signal representation with a clear interpretation as components of decreasing effect size. In contrast to other factorisations that deploy biophysical constraints, e.g. to enforce non-negativity in both weights and basis functions (Christiaens et al., 2017; Jeurissen et al., 2015; Reisert et al., 2014), the SVD deliberately avoids any such constraints. As such, we can ensure that the optimised multi-shell HARDI scheme is driven by the data itself, not by implicit assumptions of a particular tissue model.

The optimisation suggests that the optimal number of directions per shell can be expressed as a fraction of the total number of volumes to be acquired. Furthermore, the contrast to noise ratio of the estimated coefficients will scale with the square root of the total number of measurements. These results can therefore easily be tailored to the particular circumstances under which the protocol is to be used, using the recommendations outlined below:

1. Identify the desired imaging parameters (resolution, spatial coverage, etc.), which will determine the SNR and repetition time of the sequence. Divide the scan time allocated to this sequence by the TR to obtain an estimate of the total number of volumes to be acquired;
2. based on the SNR and total number of volumes, compute the CNR for the various coefficients for the 2, 3 & 4 shell scenarios, using equation (15);
3. decide on the number of shells that will be used, by selecting that which predicts usable CNR for all estimated coefficients (i.e. the CNR is above the noise floor for all coefficients);
4. using our previous orientation-domain approach (Tournier et al., 2013), verify that the number of directions for each *b*-value is sufficient to avoid aliasing, given the noise level in the images;
5. if the number of directions per shell is not sufficient to avoid aliasing in the angular domain, the data acquisition will not provide optimal characterisation of the dMRI signal. There are several ways to address this issue:

- Increase the number of volumes in the acquisition, and hence the scan time, to meet the minimum sampling requirements.
- Select a lower number of shells, as this increases the number of directions for the remaining shells. This decision will depend on whether the error introduced by an inadequate *b*-value dependence model (i.e. having too few shells) is greater than that introduced by an inadequate angular dependence model (i.e. having too few directions).
- Redistribute directions between shells to ensure the minimum sampling requirements are met, striving to maintain the number of directions per shell as close to those identified as optimal. While this implies a departure from the optimal parameters for overall CNR, the actual impact is likely to remain acceptable for small adjustments, and can be evaluated using the approach outlined in this study.

It is interesting to note that the *b*-values identified as optimal are very robust across subjects (Figure 8): for the 3-shell case, *b* = 400 s/mm^2^ and 1000 s/mm^2^ are identified in all subjects with high reproducibility (range is *b* = 397 – 411 s/mm^2^ and *b* = 988 – 1001 s/mm^2^ respectively). There is more variation for the highest *b*-value (ranging from *b* = 2,360 to 3,700 s/mm^2^), as can be seen from Figure 8. Moreover, these values are relatively insensitive to changes in *T*_2_ (Figure 7), and remarkably consistent when different numbers of shells are selected (*b*=400 s/mm^2^ is present in all schemes, *b*=1000s/mm^2^ is present for the 3 & 4 shell schemes) (Figure 6). However, given the smoothness of the principal components, minor variations in the exact *b*-values used are unlikely to have a large impact on the optimality of the acquisition.

Being entirely data-driven, the approach inherently provides parameters optimal for the particular cohort under investigation. In the present work, we focus on the neonatal age range, and the parameters identified should therefore only be considered optimal for this age range specifically. This approach could also be used to optimise protocols for ex vivo scanning, for instance. However, we note that the *b*-value optimisation procedure used here did not explicitly take the orientation information into account (beyond ensuring the minimum sampling density requirements are met); this approach may yield sub-optimal results if applied to the adult case where the angular dependence is much stronger. In contrast, the neonatal brain exhibits much lower anisotropy, and many of the most interesting features in this age range are likely to manifest as changes in microstructure rather than fibre orientation; such features are likely to be better characterised by focusing on the *b*-value dependence of the mean DW signal (e.g. (Kaden et al., 2015; Reisert et al., 2017)), motivating in part the current approach. Joint optimisation of both angular and *b*-value dependences requires extended orthonormal *q*-space decompositions (Christiaens et al., 2019), and is the subject of ongoing work.

As shown in Table 3, the lowest *b*-value selected by the algorithm, at b≈400s/mm^2^, is lower than has typically been used in previous multi-shell acquisitions (notably the Human Connectome Project), although in line with recommendations for diffusion kurtosis imaging (DKI) (Jensen et al., 2005; Poot et al., 2010) and neurite orientation density and dispersion imaging (NODDI) (Zhang et al., 2012). Furthermore, lower *b*-values are expected considering that mean diffusivity is higher in neonates than in the adult brain. On the other hand, the highest *b*-value selected is similar to those identified as optimal for fibre orientation estimation in adults (Alexander and Barker, 2005; Tournier et al., 2013); the acquisition scheme should therefore be well suited to tractography applications. Finally, the number of DW directions per shell increases with higher *b*-value, most likely due to the stronger signal attenuation (coincidentally, increasing sampling density is also required to characterise the increasingly complex angular dependence of the signal at higher *b*-values).

A question that naturally arises from this work is whether and how the sampling directions should be optimised across shells. It has been suggested in previous work that the sampling directions should be chosen to provide uniform angular coverage across all shells (Caruyer et al., 2013). In contrast, in this study the directions were optimised for uniformity across each shell independently, with no attempt to introduce any interaction between shells. We motivate our approach using sampling theory, based on similar arguments to our previous work (Tournier et al., 2013), as follows. We assume the diffusion signal is band-limited in the angular domain; i.e. the diffusion signal varies smoothly as a function of orientation, and its angular frequency spectrum contains low harmonic terms only (depending on the *b*-value). By construction, our sampling scheme is sufficiently dense that all angular frequency components have been fully captured. When these conditions are met, there is no additional information to be gained by introducing inter-dependencies across shells, since the signal is already fully sampled. Furthermore, any attempt at introducing cross-shell dependencies can only push the sampling scheme away from per-shell uniformity, and hence compromise its overall quality. We therefore feel justified to optimise the directions on a purely per-shell basis (we do however note that optimisation across shells may be beneficial in the undersampled case, where each shell is not sufficiently sampled in isolation to capture all features of the signal – in this case, a suitable model of the signal may be able to make use of the complementary orientational information provided by the different shells).

Identifying the optimal number of shells to be included in the final protocol necessarily requires additional factors to be considered, most notably *N*_*total*_, which is in turn dependent on details of the acquisition and the total feasible acquisition time. The minimum angular sampling requirement imposes an upper limit on the number of shells that can be acquired for a chosen *N*_*total*_ since increasing the number of shells necessarily reduces the number of directions per shell. However, these requirements are easily met for large *N*_*total*_, as is the case for the dHCP. It is notable that adding higher shells can strongly suppress the number of samples available for lower shells as, at least in the neonatal case, almost 50% of the samples must be obtained using the maximum *b*-value. We used the per-voxel maps of weights for each component to help inform this balance, based on an expectation that meaningful components should exhibit anatomically plausible spatial patterns that are consistent across subjects. As shown in Figure 5, the maps for the 4^th^ component do exhibit some anatomically plausible contrast (notably in the posterior limb of the internal capsule), whereas the 5^th^ component is much less consistent across the test subjects, suggesting it was strongly impacted by noise and artefacts, probably related to residual misregistration. For this reason, and given the non-negligible drop in CNR for all components when increasing the number of shells (Table 4), we decided to use a 3-shell (+ *b*=0) scheme for the final dHCP protocol, with 300 volumes acquired within 20 minutes using multiband factor 4 (Hutter et al., 2017).

## Conclusion

The approach presented in this study provides a means to design an optimal acquisition multi-shell HARDI protocol, based on maximising the information content of the data, independently of any particular reconstruction method. It is based on empirically measured data, with no reliance on any assumed model of microstructure. It focuses primarily on the mean DW signal per shell, with separately-derived safeguards to ensure adequate angular sampling; this is appropriate for neonatal data given the low angular contrast in this age range. The analysis performed here suggests that a *b*=0 + 3 shells acquisition is suitable for neonatal imaging, consisting of *b =* 0, 400, 1000, 2600 s/mm^2^ with numbers of DW directions per shell in proportions 6:19:28:47; these were employed in the final diffusion MRI protocol for the dHCP project.

## Acknowledgements

This work received funding from the European Research Council under the European Union’s Seventh Framework Programme (FP7/2007-2013)/ERC grant agreement n° 319456, and was supported by the Wellcome EPSRC Centre for Medical Engineering at King’s College London (WT 203148/Z/16/Z), and by the National Institute for Health Research (NIHR) Biomedical Research Centre based at Guy’s and St Thomas’ NHS Foundation Trust and King’s College London. The views expressed are those of the authors and not necessarily those of the NHS, the NIHR or the Department of Health.

## Appendix A: Optimising against DTI indices

To illustrate the issues inherent in using specific model parameters to optimise general acquisition parameters, we performed a simple analysis based on the diffusion tensor model, geared towards identifying the optimal subset of *b*-value shells from the full available set, an approach similar to that used in a recent study (Howell et al., 2018). This analysis relies on the availability of an oversampled acquisition that can serve as the ‘reference’, and from which candidate acquisitions can be formed by taking subsets of the data. It is premised on the notion that the best ‘reference’ reconstruction is obtained by taking all available data into account. To be deemed optimal, a candidate acquisition should provide results that closely match these reference results. Here, we investigate all possible 2-shell subsets of the reference data (all *b*=0 volumes were included in all candidates), as listed in Figure 1. All candidate schemes therefore consist of the same amount of data: 5 *b*=0 volumes + 2 × 50 diffusion-weighted volumes.

The data collected as part of this study provides an oversampled acquisition suitable for this type of analysis. To explore the issues with such an approach, we tested three different commonly-used diffusion tensor fitting algorithms: an ordinary least-squares fit to the log-signal (OLS); a weighted least-squares fit to the log-signal (WLS); and an iteratively re-weighted least-squares fit to the log-signal (IWLS) (Basser, 1995; Koay et al., 2006; Salvador et al., 2005; Veraart et al., 2013). For each fitting approach, the reference reconstruction and all candidate reconstructions were computed using that approach, and the results of each candidate acquisition were always compared only to the reference computed using the same fitting algorithm.

As in (Howell et al., 2018), we look at FA and MD as the outcome metrics of interest, and quantify the discrepancy between each candidate acquisition and the reference by computing the absolute difference in the computed metrics, normalised to the corresponding reference value. The median absolute difference is computed within a conservative brain mask for each subject, and subsequently averaged across subjects. The resulting discrepancy values are shown in Figure 1, for FA & MD (columns) and all three fitting algorithms (rows), for all candidate diffusion schemes (*x*-axis). The OLS results differ markedly from the other fitting approaches, with more minor differences between WLS and IWLS. More importantly, the acquisition identified as ‘optimal’ differs depending on which fitting approach and output metric was used.

In addition, we also computed the discrepancy between the reference results computed using the different fitting approaches. For any two of the fitting approaches, the absolute difference in the reference values produced (i.e. obtained using all available data) were computed, and the median calculated. These results are shown in the bottom right panel of Figure 1, for both FA and MD metrics, for all pairs of fitting methods. Note that the differences observed in this case are as great or larger than those observed in the previous analysis. In other words, the estimator used to compute the metrics has a greater impact on the results than the combination of *b*-values used in the acquisition. Furthermore, in the absence of ground truth, there is no principled way to decide which fitting algorithm should be used.

As can readily be appreciated, there is little consistency in these results, making it near-impossible to identify which of these schemes can legitimately be claimed to be optimal in any sense. These results would no doubt differ again if other higher-order models had been included in the comparison, including, but not limited to: Q-ball imaging (Descoteaux et al., 2007; Hess et al., 2006; Tuch, 2004), ball & sticks (Behrens et al., 2007), NODDI (Zhang et al., 2012), and/or spherical deconvolution (Tournier et al., 2007; Jeurissen et al., 2014; Christiaens et al., 2017). Moreover, most of these models can also be fitted using different approaches, each with different user-adjustable parameters (e.g. amount of regularisation, harmonic order, etc), and produce different types of output metrics. With such a wide range of models, parameters and outcomes, it is clear that a different approach is required, motivating the current study.

